# West Nile virus in crocodiles and mosquitoes in Zimbabwe

**DOI:** 10.1101/2021.02.19.431612

**Authors:** Marlène Roy, Fortunate Mufunda, Deirdre Slawski, Charmaine Mutswiri, Irene Mvere, Katherine Slawski, Pamela Kelly, Ilaria Piras, Annemarie Bezuidenhout, Tapiwanashe Hanyire, Bryan Markey, Gerald Barry

**Author notes:** **Address for correspondence**: Gerald Barry, Veterinary Biosciences, School of Veterinary Medicine, University College Dublin, Belfield, Dublin 4, Ireland.

## Abstract

We detected, for the first time, West Nile virus lineages 1 and 2 in Zimbabwe in mosquitoes and crocodile tissue samples, including fluid from egg waste. Our results provide evidence of WNV circulation in Zimbabwe, suggesting that an evaluation of the risk to humans and susceptible animals should be considered.

## Text

The flavivirus, West Nile virus (WNV), is spread primarily by *Culex* mosquitoes among birds and many spillover hosts, including humans, horses and crocodiles (1)

There are 9 lineages of WNV, but lineages 1 and 2 are the most commonly associated with human outbreaks of disease (1). Lineage 1 is typically found in Europe, Africa, Asia, the Americas and Australia. Lineage 2 strains are found in Africa and Europe (2). While WNV is an arbovirus, a number of studies have highlighted the potential for both mosquito and water-borne transmission (3,4)

There is a paucity of information on the viruses that may be present in Northern Zimbabwe. In collaboration with a Nile crocodile (*Crocodylus niloticus*) farm we collected post-mortem tissue samples including skin, blood, kidney, brain, lung and liver from crocodiles that had been culled as part of routine farming practices, along with fluid from egg waste. All samples were collected with the approval of the UCD Animal Research Ethics Committee; no procedures were conducted on living animals and no animal was euthanized for the sole purposes of the study. Pooled samples of mosquitoes and midges caught on and off farm were also collected, using light traps. The traps contained mosquitoes including members of the genera *Anopheles*, *Aedes* and *Culex* as well as midges belonging to the *Chironomidae, Cecidomyiidae* and *Ceratopogonidae* families.

Total RNA was extracted from 16 insect pools (6 mosquito and midge mixed pools, 8 mosquito only pools, 2 midge only pools) and 180 crocodile tissue samples by a Trizol (Thermo Fisher Scientific) method according to the manufacturer’s instructions. Reverse transcription (RT) was performed using SuperScript III Master Mix (Thermo Fisher Scientific) and PCR was performed using the 5x HOT FIREPol SolisGreen PCR Mix (Solis BioDyne). The cDNA was checked using the housekeeping gene 18S ribosomal RNA (5’ AGG ATC CAT TGG AGG GCA AGT 3’ and 5’ TCC AAC TAC GAG CTT TTT AAC TGC A 3’) (5) followed by PCR screens using the pan-flavivirus primer set: Flav-fAAR 5’ TAC AAC ATG ATG GGA AAG AGA GAG AAR AA 3’ and PFlavrKR 5’ GTG TCC CAK CCR GCT GTG TCA TC 3’ (6). A 265 bp DNA fragment of the NS5 gene was amplified from mosquito only, mosquito/midge and crocodile tissue pools but not from midge only pools. Virus replication was confirmed in select insect and crocodile samples through RT reactions using the pan-flavivirus primer sets in both sense and antisense orientations, followed by end-point PCR.

A selection of pan-flavivirus positive samples were then tested specifically for the presence of WNV RNA using primers targeting the WNV capsid gene (130 bp fragment; 5’ GCCGGGCTGTCAATATGCTAAAA 3’ (7) and 5’ AAGAACGCCAAGAGAGCCAAC 3’ (8)). All 31 pan-flavivirus positive samples tested were positive for WNV.

Further confirmation was provided by lineage specific PCR on 22 of these 31 samples. A WNV lineage 1 primer set: 5’ TGCCTAGTGTCAAGATGGGG 3’ and 5’ ACTCTTCCGGCTGTCAATCA 3’, was designed (reference sequence NC_009942.1) to amplify a 200 bp fragment of the NS3 gene while a WNV lineage 2 (NC_001563.2) primer set: 5’ AACTGATCATGAAGGACGGC 3’ and 5’ ACATCTGCGCGTATGACTTC 3’, was designed to amplify a 141 bp fragment of the RNA polymerase.

Of the 22 samples tested, 8 were positive for lineage 1 and lineage 2, 2 were positive for lineage 1 only while 12 were positive for lineage 2 only. Sanger sequencing was carried out on the positive PCR results (Table 1). Unfortunately, the quality of samples was not sufficient in several cases leading to sequences that were not usable. BLAST analysis of the good quality sequences revealed that three mosquito/midge pools, a crocodile egg waste fluid pool and two crocodile lung/liver samples showed high similarity (at least 92%) to sequences from WNV lineage 2 isolates including, in 2 cases, WNV strain B956 (GenBank accession no. AY532665), the first WNV strain identified from a human patient in Uganda in 1937 (9). Two insect samples also gave a very short sequence matching WNV lineage 1.

To the best of our knowledge, no studies in Zimbabwe have reported the presence of WNV in crocodiles, insects or humans. WNV has been detected in birds (barn swallow and brown-throated martin) by RT-PCR in central Zimbabwe (10). We report here the detection of WNV RNA for the first time in northern Zimbabwe in mosquitoes and crocodile samples, highlighting the possibility of local transmission. Our results suggest that WNV transmission among crocodiles may involve vector-borne infections, but other transmission routes could also be important. In particular, the finding of WNV RNA in fluid from egg waste suggests the possibility of vertical transmission. This requires further investigation as it could have significant implications for the prevention of WNV introduction into farms and zoos. This report highlights potential veterinary and human health concerns and improved surveillance of this and other potential pathogens in the area is warranted.

## Acknowledgments

The support of Gary Sharp, Chief Executive Officer of Padenga Holdings Limited is gratefully acknowledged. This research was approved by the Research Council of Zimbabwe. This work was supported financially by Padenga Holdings Limited.

## Author Bio

Dr. Roy is a post-doctoral researcher working in the research group of Dr. Barry, at the School of Veterinary Medicine, University College Dublin, Ireland. Her research interests include the molecular study of arboviral diseases and virus-host interactions.

---

**Table.**
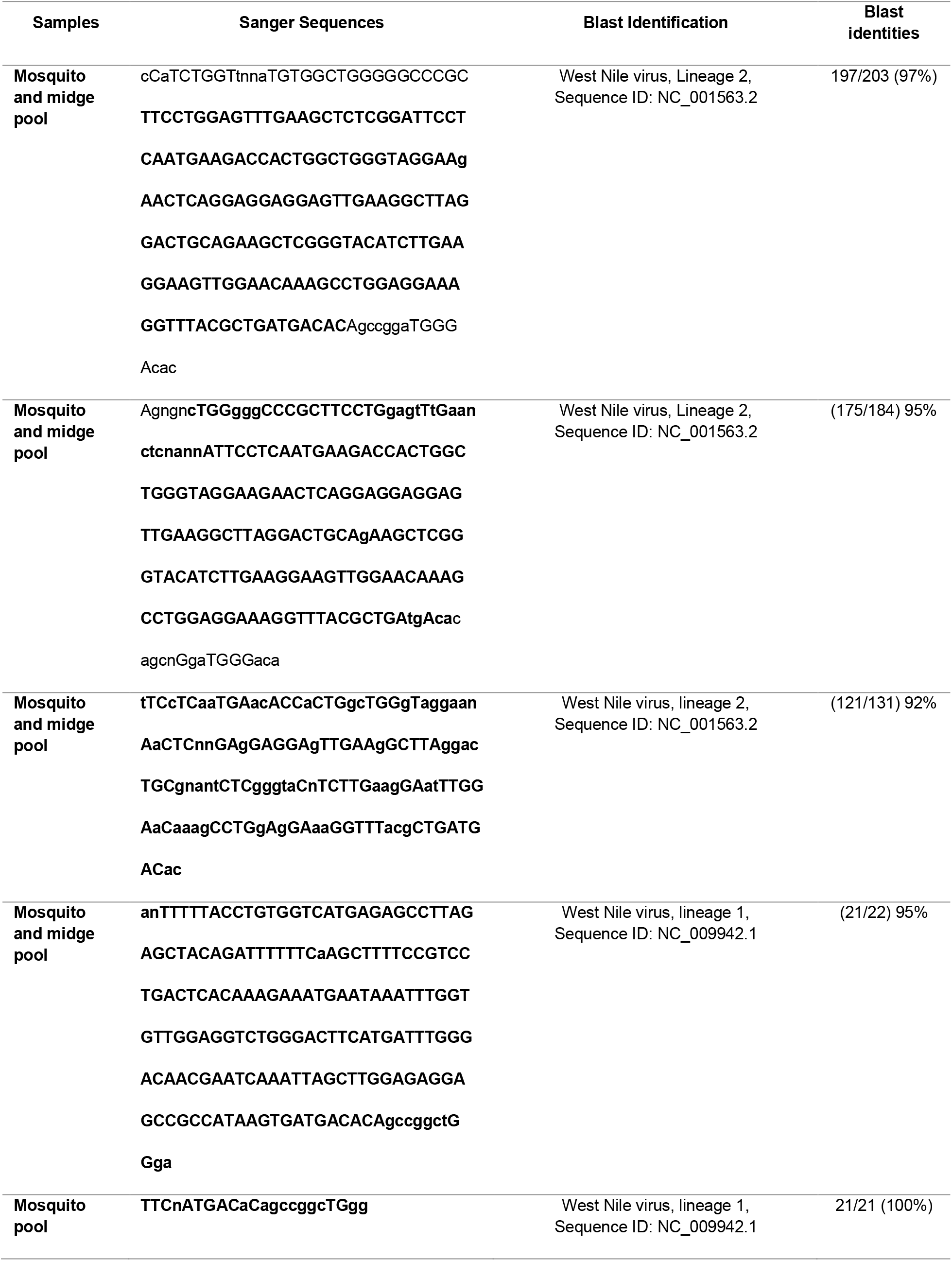
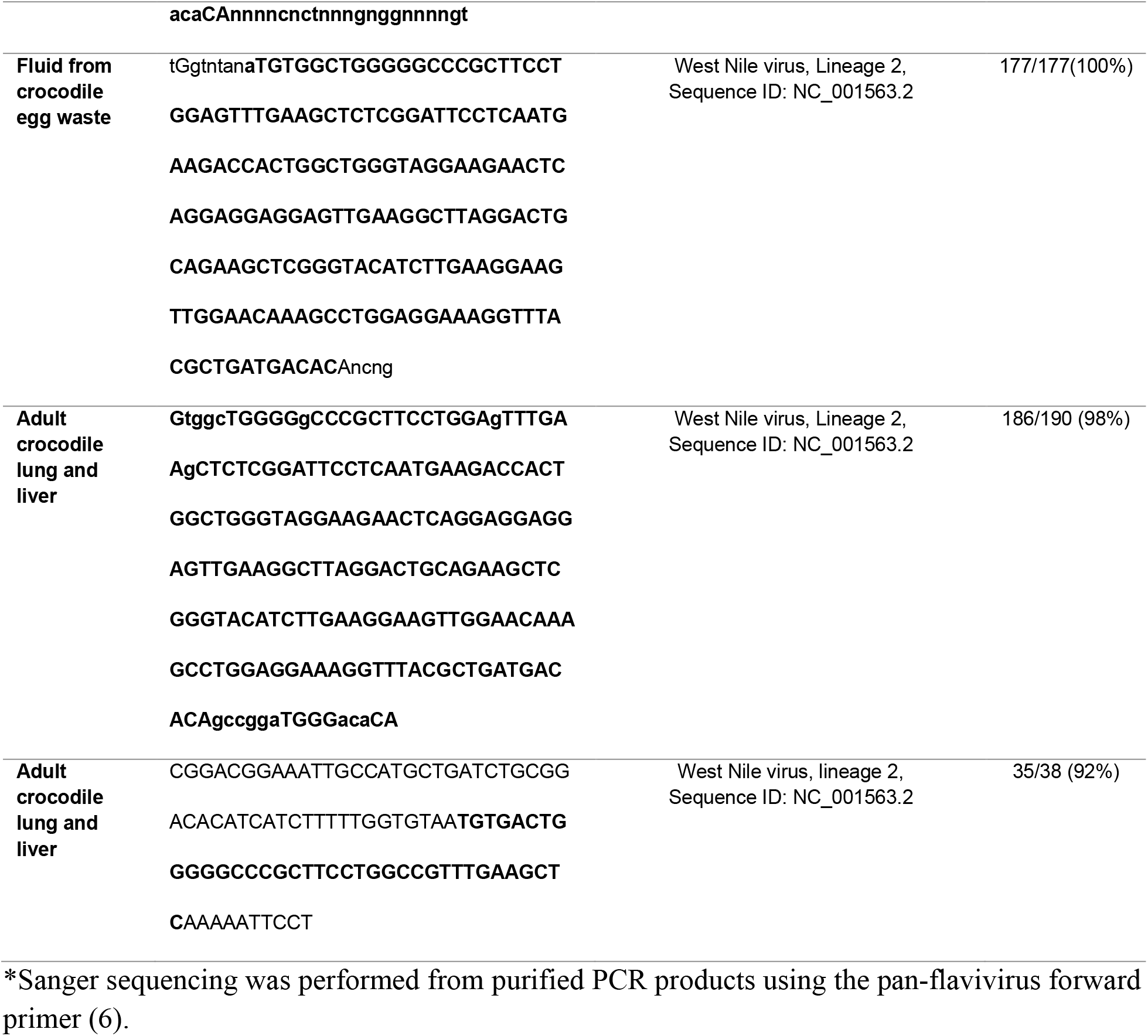
WNV sequences from insect and crocodile tissue samples*.

